# Island invasions by the non-native vinegar fly *Drosophila suzukii* and its parasitoid wasps

**DOI:** 10.1101/2025.09.04.674335

**Authors:** Paul K. Abram, Elizabeth H. Beers, Charlie C. Coslor, Benjamin Diehl, Michelle T. Franklin, Louis B. Nottingham, Jason Thiessen, Matthew Tsuruda, Juli Carrillo

## Abstract

Understanding how island characteristics influence the establishment and impact of invasive species and their natural enemies could inform both island biogeography and biological control theory. We studied the occurrence and relative abundances of the globally invasive fruit fly, *Drosophila suzukii*, and its recently introduced larval parasitoids, *Leptopilina japonica* and *Ganaspis kimorum*, across islands with varying sizes and levels of human-mediated transport in the Gulf and San Juan Islands of British Columbia (Canada) and Washington State (USA). Across two years and 58 sites, we collected *D. suzukii* and its parasitoids from wild blackberry, *Rubus armeniacus*, fruit. We predicted that parasitoids were more likely to be present on larger islands with higher levels of human activity and higher *D. suzukii* densities, and that the less specialized parasitoid species (*L. japonica*) would be more likely to establish on islands. We detected *D. suzukii* across all islands, indicating widespread establishment of this invasive pest. In contrast, we observed parasitoids on fewer than half of the islands. *Leptopilina japonica* was the only parasitoid of *D. suzukii* detected on islands. Parasitoid presence was marginally positively associated with island area and host density, but not average annual vehicle-ferry traffic (an indicator of human-mediated propagule pressure). Parasitism levels were low throughout the study region and we did not observe lower relative abundances of *D. suzukii* on islands where parasitoids were present – in fact, the relative abundance of *D. suzukii* tended to be higher on islands where the parasitoid *L. japonica* was detected. These findings suggest that island characteristics, host density, and a consumer’s host specificity may be associated with early establishment of introduced consumers on islands, but that a consumer’s presence may not inevitably result in host population suppression, at least over the short term.

## Introduction

Island biogeography theory has been a central framework for explaining patterns of occupancy, abundance, and community composition of organisms on islands of different sizes and degrees of isolation (MacArthur and Wilson 1967; Losos and Ricklefs 2009). Dispersing species typically establish earlier on larger, less isolated islands, leading to increased species richness compared to smaller, more isolated islands. Distance from mainland source areas was historically considered the main determinant of island isolation (MacArthur and Wilson 1967); however, variation in human activity that leads to variation in propagule pressure (i.e., the number and size of introduction events of a given species) may now be a stronger predictor of organisms’ dispersal to islands (Helmus et al. 2014). While the main focus of island biogeography theory is explaining differences in the number of species occupying different islands, it may also be useful for understanding the susceptibility of different islands to contemporary invasions by novel species and the strength and nature of resulting trophic interactions among them (Holt 2009).

A species’ trophic position and level of specialization are important determinants of its ability to establish on islands of different sizes (Holt et al. 1999; Holt 2009; Gravel et al. 2011). In particular, island size is thought to have a stronger effect on a consumer than its resources (the “trophic rank hypothesis”; Holt et al. 1999). This is thought to be because larger islands have a greater diversity and quantity of resources to support consumers (Schoener 1989; Holt 1999; Holt 2009) and are more likely to provide the necessary geographic area needed to stabilize consumer-resource interactions (Hassell et al. 1991; Hassell 2000). Consumers, if they are at lower regional abundance than their resource, may also be disproportionately more likely to occur on large islands because of simple numerical effects (Srivastava et al. 2008). Specialized consumer species are generally more sensitive to island area than generalist species because their persistence relies on withstanding the demographic stochasticity of a smaller number of host species (Freeland 1983; Steffan-Dewenter and Tscharntke 2000; Kruess and Tscharntke 2000). Thus, on islands of a given size, consumers are likely to establish less often than their resource species, with establishment becoming less likely as their level of specialization increases.

The arrival of consumers on islands may, in turn, shape how island size affects the abundance of lower trophic levels that they consume (Schoener and Spiller 1996; Schoener et al. 2016; Piovia-Scott et al. 2017) (Figure 1). If the resource species initially establishes on islands in the absence of specialized consumer species that are present in source (mainland) areas and exert strong top-down control, the resource species may experience ‘enemy release’ (Keane and Crawley 2002) on islands and reach higher abundances than on the mainland (Figure 1). On islands, in the absence of their natural enemies, we might expect resource species to have higher population densities on larger islands. Contrary to the implicit assumptions of island biogeography theory, which predicts flat area-density relationships, empirical evidence shows that in general, some groups of animals (including insects) tend to have higher population densities on larger islands (Connor et al. 2000). This is consistent with the resource concentration hypothesis (Root 1973) which states – in line with often observed empirical evidence – that larger resource patches have higher densities of consumers, possibly due to movement behaviour of consumers (herbivores are more likely to find and remain in large patches) (Connor et al. 2000). If consumers are then more likely to initially establish on larger islands, their subsequent suppression of resource populations on those large islands could lead to a decrease in the slope of the relationship between island area and the density of the resource species over time (Figure 1). We term this the “trophic dampening” hypothesis, referring to the dampening effect that parasitism could have on the island area-consumer density relationship. Whether this might occur would depend, in part, on the strength of resource population suppression by consumers.

**Figure 1.**
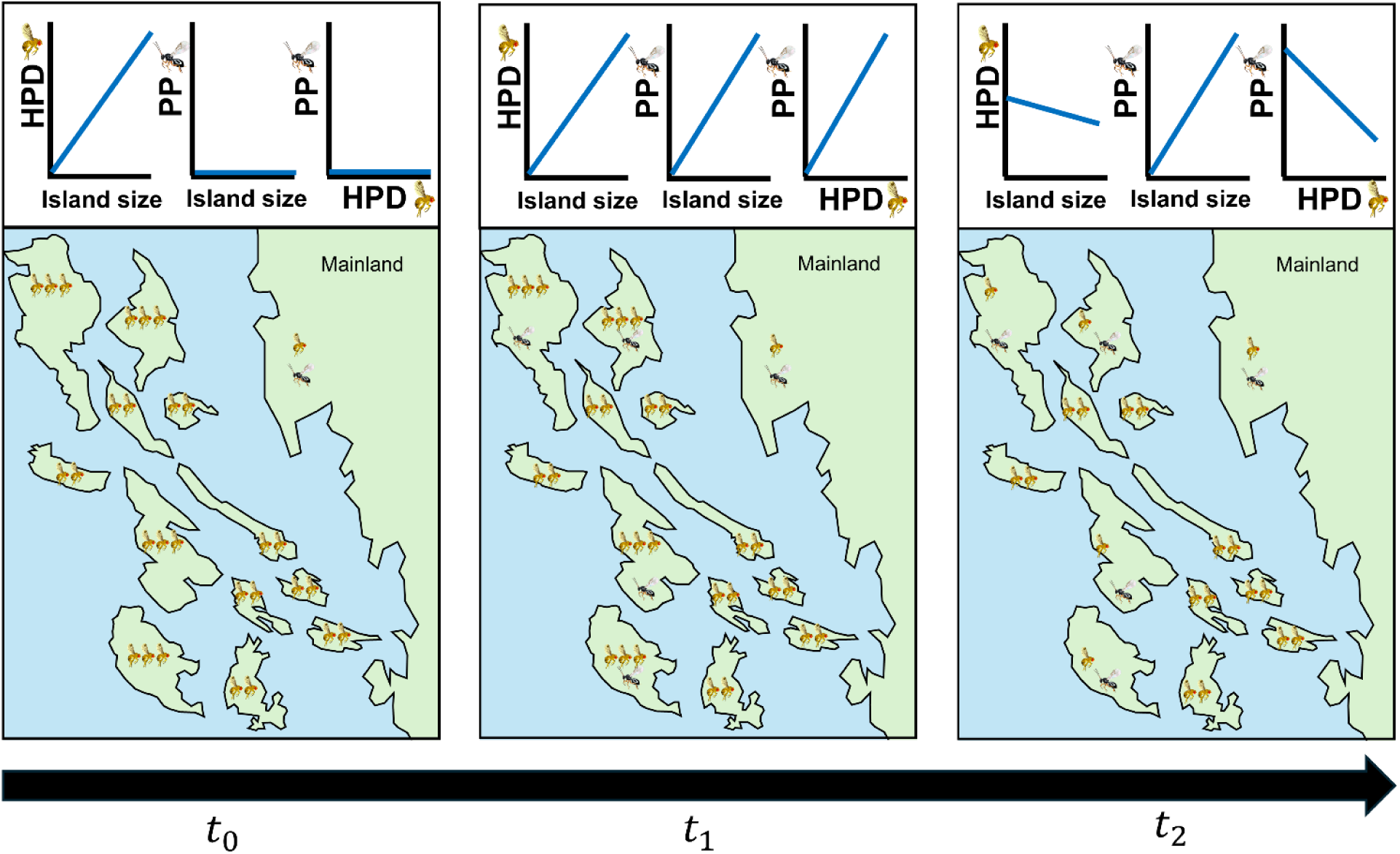
A hypothetical framework for how host population densities (HPD) and parasitoid presence (PP) are predicted to be associated with island size at different invasion stages of a host (fly) and its parasitoid (wasp), assuming widespread establishment of the host and strong population suppression of the host by the parasitoid. At t_0_, where the host has invaded all islands off the mainland but the parasitoid has not, the host reaches larger mean population densities (indicated by a greater number of flies) on islands, but especially higher on larger islands if enemy release is more important than abundance-area relationships. At t_1_, the parasitoid begins to invade islands and is more likely to establish on larger islands which have higher host population densities that act as receptive bridgeheads. At t_2_, the parasitoid has suppressed the host populations on larger islands, the slope of the relationship between host population density and island size decreases and the presence of parasitoids becomes negatively associated with host population densities (“trophic dampening”).

Many island ecosystems have been invaded by non-native species at multiple trophic levels (Moser et al. 2018), including some intentional introductions of consumers for biological control (Heimpel and Mills 2017). Islands have played an important role in the history of biological control, with some efforts resulting in strong pest suppression and others causing negative ecological consequences (Heimpel and Mills 2017). In some analyses, biological control agents have established more readily on islands than mainland areas (reviewed in Heimpel and Mills 2017), yet it remains unclear whether pest suppression is consistently stronger on islands or how it is influenced by island characteristics (Wyckhuys et al. 2022). Because islands offer semi-independent ecological units that differ in size and isolation, they provide promising but under-utilized systems for testing ecological theory and biocontrol outcomes (Wyckhuys et al. 2022). In particular, mainland-to-island spread of biological control agents could help to understand how biological control agent traits, such as trophic specialization, interact with island features to influence both establishment and pest population densities. While prior studies have examined parasitoid-host and predator-prey interactions on small or remote islands (e.g., van Nouhuys and Hanski 2002; van Nouhuys and Lei 2004; Schoener et al. 2016), less is known about such dynamics on larger, more human-impacted islands.

Here, we conducted a two-year study of the occurrence and relative abundance of an invasive pest, spotted-wing Drosophila, *Drosophila suzukii* (Matsumura) (Diptera: Drosophilidae), and two species of introduced parasitoid wasps in an island landscape highly shaped by anthropogenic activity and transport: the Gulf and San Juan Islands off the coast of British Columbia (Canada) and Washington State (USA). The two parasitoid wasp species, *Leptopilina japonica* Novkovic & Kimura and *Ganaspis kimorum* Buffington (Hymenoptera: Figitidae), were under consideration for intentional biological control introductions against *D. suzukii* in the USA, Canada and Europe when both species were found to already be established, common and widespread in mainland areas of British Columbia and Washington State, as well as Vancouver Island (Abram et al. 2020, 2022a; Beers et al. 2022). Whether or not *D. suzukii* (first detected in the mainland areas in 2008-2009) and its parasitoids (first found in 2016-2019) had spread to more remote areas, including the network of much smaller islands nestled between Vancouver Island and the Mainland (Figure 2), was unknown. Due to a lack of long-term longitudinal data from the mainland, the impact of the parasitoids on *D. suzukii* populations in their introduced range are still unclear. The recent arrival of *D. suzukii* and its associated oligophagous (*L. japonica*) and specialized (*G. kimorum*) parasitoid species on the mainland could be thought of as a ‘natural experiment’ to examine how characteristics of adjacent islands (isolation, size) and parasitoid specialization are associated with their occupancy of different islands. It is also an opportunity to test whether parasitoid establishment on islands results in effective biological pest control, and how these outcomes are associated with island size and isolation.

**Figure 2.**
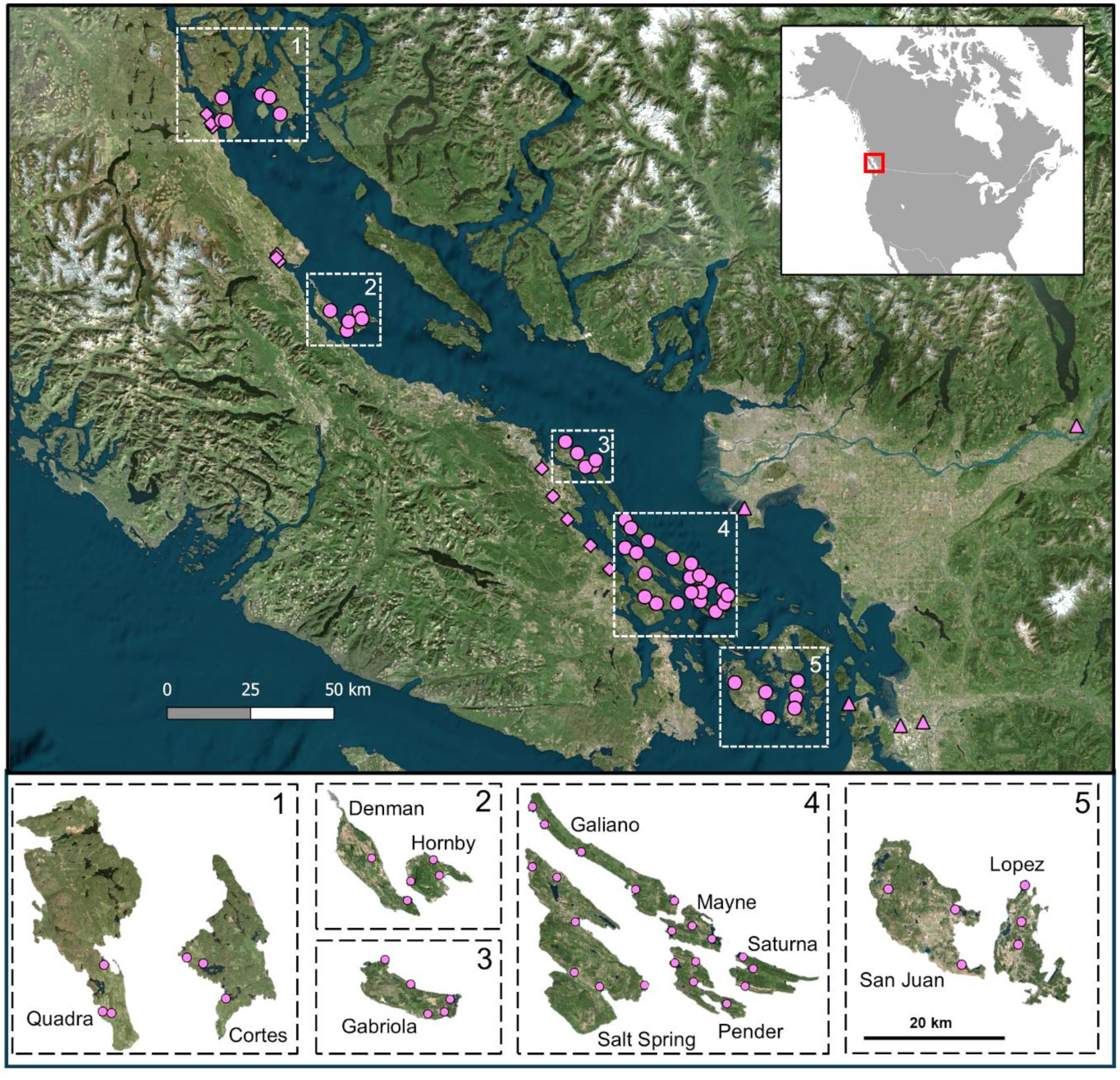
Sampling sites for *Drosophila suzukii* and its larval parasitoids in coastal British Columbia, Canada and Washington State, USA. The inset in the top right corner shows where the study region is located in North America (indicated with red box). Pink symbols in upper panel: Triangles – sampling sites on the mainland; diamonds – sampling sites on Vancouver Island; circles – sampling sites on the Gulf Islands. The bottom panel shows zoomed-in depictions of the twelve studied Gulf Islands (to scale; numbers in top right corners correspond to those inside white dotted boxes on the upper map) and the sampling sites on each.

Based on island biogeography theory, we first predicted that parasitoids would be more likely to be present on islands that are larger and less isolated. Because the island system we studied is connected by ferry services that transport hundreds of thousands of vehicles among mainland areas and the islands each year, and *D. suzukii* is associated with commodities commonly transported by humans (fruit), we reasoned that the main source of relative isolation of some islands would not be geographic distance from mainland areas; it would be lower levels of human activity. We also predicted that parasitoids would be more likely to be present on islands with higher average host densities, because following initial introduction events, parasitoids would be more likely to detect and locate hosts, and subsequently establish breeding populations, when hosts are at higher densities (*sensu* the ‘receptive bridgehead hypothesis’; Weber et al. 2021). If parasitoids are in the early phases of establishment (i.e. they are uncommon and parasitism levels are low), they might be more likely to be found on islands with higher host densities (Figure 1). In this case, if most island populations of *D. suzukii* are experiencing strong enemy release, we might expect parasitism levels to be low and *D. suzukii* populations to be at high densities on islands relative to the mainland. On the other hand, if the parasitoids are already well established and strongly suppress *D. suzukii* populations on islands where they had established (i.e. they are common and parasitism levels are high), we would expect islands where parasitoids are present to have lower *D. suzukii* population densities (i.e., number of larvae per sampled fruit) and higher parasitism levels, similar to those in mainland areas (Figure 1). Finally, all else being equal, we would expect the less specialized parasitoid species (*L. japonica*) to be more likely to establish on an island with a given set of characteristics than the more specialized parasitoid species (*G. kimorum*).

## Materials and Methods

### Study system

*Drosophila suzukii*, native to Asia, has become a major globally invasive pest of soft-skinned fruit crops since it was first detected in North America and Europe in the late 2000s (Asplen et al. 2015). This vinegar fly is able to lay its eggs and complete its larval development in the ripening fruits of many different cultivated and wild plants (both native and non-native) in agricultural, urban, semi-natural, and natural environments (Kenis et al. 2016; Abram et al. 2022a). Multiple *D. suzukii* larvae can develop in a single berry. In British Columbia, Canada, *D. suzukii* overwinters as an adult and has multiple overlapping generations over the course of its seasonal reproductive period (May to October), during which its populations reproduce on plants with different fruiting phenology and peak in density from August to September (Abram et al. 2022a). During its seasonal peak in abundance in our study area, the most widespread and common host plant that reliably harbors *D. suzukii* and its parasitoids is the non-native weedy Himalayan blackberry, *Rubus armeniacus* (Abram et al. 2022a). First introduced to North America in the late 1800s, *R. armeniacus* is now widespread and common in disturbed and riparian areas in coastal British Columbia and Washington. Based on these features, we sampled from *R. armeniacus* across all sites to estimate *D. suzukii* population density.

The parasitoid wasps *L. japonica* and *G. kimorum* lay their eggs inside the larvae of *D. suzukii* feeding within fruit (Stahl et al. 2024; Rossi Stacconi et al. 2025). They kill their host once it has formed a puparium within the fruit or the soil/leaf litter below. Native to Asia and previously identified as candidates for importation and release as biological control organisms for *D. suzukii* in the USA, Canada and Europe, the two parasitoids have different levels of specialization on *D. suzukii. L. japonica* is oligophagous; it is closely associated with *D. suzukii* but can also parasitize several other Drosophilidae species (Rossi Stacconi et al. 2025). *Ganaspis kimorum*, in contrast, appears to be highly specific to *D. suzukii* (Stahl et al. 2024). The natural dispersal abilities of these parasitoids are unknown, but it is believed that human activity is primarily responsible for recent unintentional introductions and rapid spread of *L. japonica* to several areas of North America (Abram et al. 2020; Gariepy et al. 2024) and Europe (Rossi Stacconi et al. 2025). *Ganaspis kimorum*, on the other hand, has only established adventive populations in Washington and British Columbia; intentional biological control releases of this species are being done in other areas of Canada (Ontario), the USA (several states), and Europe (Italy, Switzerland) (Gariepy et al. 2024; Stahl et al. 2024; Rossi Stacconi et al. 2025). Why adventive populations of *L. japonica* have spread more widely than *G. kimorum* without intentional introductions is not known, but differences in climatic tolerance and level of specialization have been put forward as hypotheses (Gariepy et al. 2024; Rossi Stacconi et al. 2025).

### Study area

The Gulf and San Juan Islands (hereafter, the Gulf Islands) are a network of hundreds of islands off the coast of Washington State (USA) and British Columbia (Canada) (Figure 2). This region has a temperate climate with Mediterranean-type weather patterns characterized by warm, dry summers and mild, wet winters with infrequent snow. While they contain some preserved ecosystems (e.g. the Garry oak ecosystems and Douglas fir forests), they are subject to considerable levels of human disturbance including commercial and residential development, logging, agriculture, and tourism. They are connected to the mainland and each other primarily through provincial and state-run vehicle ferry boat services, in addition to small passenger watercraft and airplanes. Many of the Gulf Islands are located in between mainland North America and Vancouver Island, a very large (∼31,000 km^2^) and highly-populated (>860,000 people) island which, for the purposes of this study, we treated as a ‘second mainland’ area. The twelve islands that were part of this study (San Juan, Lopez, Saturna, Gabriola, Pender, Cortes, Hornby, Salt Spring, Mayne, Quadra, Denman, Galiano) have areas of 22 to 310 km^2^, year-round human populations ranging from 465 to 11,798 residents, and receive an average of between 17,342 and 351,624 passenger and commercial vehicles from ferry services annually (2018-2023) (Statistics Canada 2025; BC Ferries 2024; Washington State Department of Transportation 2024).

### Sampling to determine pest and parasitoid occurrence and relative abundance

In 2022 and 2023, we sampled a total of 10,440 *R. armeniacus* fruit from 58 sites: 5 sites on mainland British Columbia and Washington, 10 sites on Vancouver Island, and 2–6 sites on each of the 12 Gulf islands (43 sites total on these islands). We visited two additional sites on Vancouver Island (Port Hardy, Campbell River) but were missing some samples (because not enough ripe berries were present on at least one visit) and so these sites were excluded from the analysis. We selected sites based on accessibility (e.g. bordering roadsides, parking lots, public parks and beaches) and, when logistically possible for island sites, were selected to spread out sampling locations to different areas of each island. We did not have the resources to do detailed vegetation surveys of the sampling areas, and there are no reliable, publicly available data regarding *R. armeniacus* distribution on the Gulf Islands. However, there are data available on broader-classification land cover types (CEC 2023). Forests are the dominant habitat type on all of the islands, with the rest of each island’s area made up mostly of cropland (mostly forage crops), shrubland, grasslands, wetlands, and urban areas (Appendix S1: Figure S1). All of these land cover types would be expected to potentially host *R. armeniacus*, *D. suzukii*, and the two parasitoid species (P. Abram, personal observations).

We visited each site twice per year; once in early to mid-August (range of sampling dates: August 8–22) and once between late August and mid-September (August 30-September 14). We selected sampling dates based on known or predicted phenology of berry ripening in different areas of the sampling region. We accounted for the possible effect of variation in sampling timing on *D. suzukii* abundance in the analysis (see *Statistical analyses*). On each site visit, we sampled 45 berries haphazardly from available fruit (ripe without visible external damage or rot) and divided into three ventilated sample containers (15 berries each) as per established sampling standards for *D. suzukii* and its parasitoids (see Abram et al. 2022b for details). While there can be some within-site variability in *R. armeniacus* berry size, based on previous research (Abram et al. 2022a), we did not expect significant among-site variability in berry size to meaningfully influence results and did not weigh berry samples.

Sample containers were transported and cooled or insulated to protect them from heat while being transported to laboratories where they were incubated at 21–23°C, under 16:8 light:dark cycles, and 40-60% RH. *Drosophila suzukii* puparia that formed in the berries were gently removed with fine paintbrushes 6-8 and 8-10 days after berry incubation, counted, and incubated under the same conditions in petri dishes with moistened filter paper; subsequent *D. suzukii* and parasitoid emergence was then counted. We initially performed this removal of puparia in cohorts with the intent of calculating more accurate percent parasitism measurements that exclude non-susceptible host life stages (Abram et al. 2022b), although after the study it was discovered that this method can potentially miss some parasitized individuals if parasitized larvae take longer to develop (J. Carrillo, unpublished data). Fortunately, these individuals would have been captured by our methods as we further incubated for at least 40 days to allow any additional *D. suzukii* and parasitoids to emerge, with insects removed within 24-72h of emergence. We calculated the total number of *D. suzukii* in each sample as the number of puparia that were picked from the sample, plus the number of adult flies that emerged from the remaining berry material. We considered this as an approximation of *D. suzukii* density at each site. We calculated the total number of parasitoids as the number of parasitoids emerging from extracted puparia plus the number emerging from the remaining berry material. Parasitism levels were too low (see Results) to justify the planned analysis of cohort-based parasitism levels, and so these analyses were not done. Fly puparia, adult flies, and adult parasitoids were identified based on the keys of Abram et al. (2022b). No Drosophilidae species other than *D. suzukii* emerged from berry material. The only two parasitoid species to emerge were *L. japonica* and *G. kimorum*. Voucher specimens are stored at the Agassiz Research and Development Centre (Agassiz, British Columbia, Canada). We pooled the three subsamples collected on the same day at each site (see above) for all analyses, so that each sample consisted of 45 *R. armeniacus* berries.

### Statistical analyses

We conducted two sets of analyses on two different spatial scales: (i) on the regional scale to compare average *D. suzukii* densities and parasitism levels between the Gulf Islands and the two mainland areas; and (ii) on the island scale (for the Gulf Islands only) to test what per-island metrics were associated with parasitoid presence.

In our regional-scale analysis, we tested whether *D. suzukii* densities were higher and parasitism levels were lower on the Gulf Islands (where parasitoids were evidently less well established) than the other two ‘mainland’ areas sampled (Vancouver Island and Mainland British Columbia and Washington State), as would be expected if *D. suzukii* were experiencing enemy release on islands. We used a generalized linear mixed model with a negative binomial error distribution, implemented with glmmTMB (Brooks et al. 2017), to compare mean *D. suzukii* densities (total number of *D. suzukii* per sample of 45 berries) among sites in the three major regions (Mainland, Vancouver Island, Gulf/San Juan Islands). Region and year were included as fixed effects, while site nested within region was included as a random effect to account for repeated measures across time. The Julian date when each sample was taken was included as an additional continuous fixed effect represented by a non-linear spline (df=3) based on visual inspection of partial residuals plots and AIC comparisons among competing models. This term was included in the models to account for variation in sampling timing among sites that could have affected *D. suzukii* populations and parasitism levels, which both tend to increase and then decline in *R. armeniacus* berries over the course of the seasonal period during which we sampled (Abram et al. 2022a); this was confirmed by inspection of model fits and partial residuals plots. We initially considered using heat units (degree days) as a measure of time during the season, but because different areas of our study region had markedly different rates of heat unit accumulation over the sampling period (Figure S2), this was found to be an unsuitable method of correcting for variation in sampling timing. Statistical significance testing (α = 0.05) was done with Type II Wald Chi-square tests. We conducted a similar GLMM with proportion parasitism as the response variable, the same fixed and random factors, and a binomial error distribution to compare parasitism levels among the three regions over the two-year study period. For both models, we used Moran’s *I* test on the Pearson residuals from our generalized linear mixed model of parasitoid presence to assess spatial autocorrelation in model residuals, using the spdep package (Bivand and Wong 2018). We used mean latitude and longitude coordinates for the sampling sites on each island to construct a spatial neighborhood matrix based on nearest neighbors (k=16 was the smallest value yielding 1 subgraph) We calculated Moran’s *I* under a randomization null model with row-standardized spatial weights. A significant Moran’s *I* statistic would indicate spatial clustering of residuals not accounted for by the predictors included in the model.

Next we conducted island-level analyses. First, to obtain per-island estimates of mean *D. suzukii* densities among the twelve sampled Gulf islands while controlling for other variables, we used a generalized linear mixed model (negative binomial error distribution) with *D. suzukii* densities (total number of *D. suzukii* per sample of 45 berries) as the response variable and island, year, and Julian date (represented by a spline with df=3; see above) as explanatory variables. We included sampling site nested within island as a random effect to account for repeated measures across time. We performed statistical significance testing (α = 0.05) with Type II Wald Chi-square tests. We calculated estimated marginal mean *D. suzukii* density for each island across years and sampling times with the emmeans package (Lenth 2024), and tested for spatial autocorrelation in the residuals using Moran’s *I* (k=11).

Then, to assess what factors are associated with the presence of parasitoids on some islands and not others, we ran generalized linear models (GLMs) with parasitoid presence (i.e., detection of at least one parasitoid over the two-year period) as the response variable. We used Firth’s bias-reduced penalized maximum likelihood estimation to model the presence of parasitoids across islands, implemented via the logistf package in R. We used profile penalized likelihood confidence intervals and p-values for each parameter estimate (Mehta & Patel 1995; Heinze & Schemper 2002). This method reduces small-sample bias and provides finite, robust estimates where standard logistic regression results in quasi-complete or complete separation (i.e., the predictor variables perfectly predict the outcomes). The continuous factors included in the model were the size of the island (in km^2^), human activity (the average number of vehicles arriving by ferry per year, in thousands), and the average density of *D. suzukii* on each island over the two-year study period (the marginal mean density from the linear mixed model described in the previous paragraph). We verified that the probability of detecting parasitoids was not due to variation in the number of sampling sites per island by adding a continuous term (number of sampling sites) to a simplified logistic regression model containing island size and *D. suzukii* density and testing its statistical significance as described above. We used ferry traffic rather than human population size on each island as the indicator of human activity because we reasoned that the former would be more likely to correlate with propagule pressure; however, the two factors are highly correlated in any case (Figure S3).

After fitting the full island-level model including all three predictors, we used likelihood ratio tests and comparisons of model estimates (β ± SE) to investigate the potential for mediating or suppressing effects of each variable on each of the two others (e.g., whether *D. suzukii* density may mediate or suppress the effect of island size or vice versa; whether vehicle traffic may mediate or suppress the effect of island size, etc.).

Finally, to test whether the presence of parasitoids on some islands could be lowering *D. suzukii* densities, and whether this effect may depend on island area (as hypothesized in Figure 1), we fit a linear model with marginal mean *D. suzukii* density as the response variable and island area, parasitoid presence (categorical – absent or present), and their interaction as predictor variables.

We had a number of additional questions that arose during data interpretation. First, we noted a potential association between parasitoid presence and island latitude. Second, we inquired as to whether variation in island land cover, particularly percent forest cover (at the site- and whole-island levels) might be associated with *D. suzukii* abundance or parasitoid presence.

We also wanted to verify that parasitoid presence was not, in fact, associated with the degree of geographic isolation from the closest known mainland population of parasitoids. We conducted several post-hoc exploratory analyses to address these questions (Appendix S1: Methods S1; Results S1), which we will refer to in the Discussion section.

All analyses were conducted with R version 4.4.1 (R Core Team, 2024). We inspected the fitted statistical models described in all sections below using the diagnostics functions available in the DHARMa package (Hartig 2024) to ensure adequate model fit.

## Results

### Presence and density of D. suzukii and parasitoids in different regions

We reared a total of 31,169 *D. suzukii* from berry samples, with emergence from 206/232 of the pooled samples of 45 berries. *Drosophila suzukii* was present in both years at all but one of our 58 sites. *D. suzukii* densities varied by region (χ²_2_=16.02; p < 0.001); their density tended to be higher on the mainland and lower on Vancouver Island and the Gulf Islands (Figure 3). At the regional level, *D. suzukii* densities were marginally higher in 2023 than 2022 (χ²_1_=3.75; p = 0.053) and varied over time within each sampling period (χ²_3_=224.00; p < 0.0001). There was residual spatial autocorrelation in *D. suzukii* densities throughout the study area not explained by the three regions, Julian date, or year (Moran’s *I* = 0.27, p < 0.0001).

**Figure 3.**
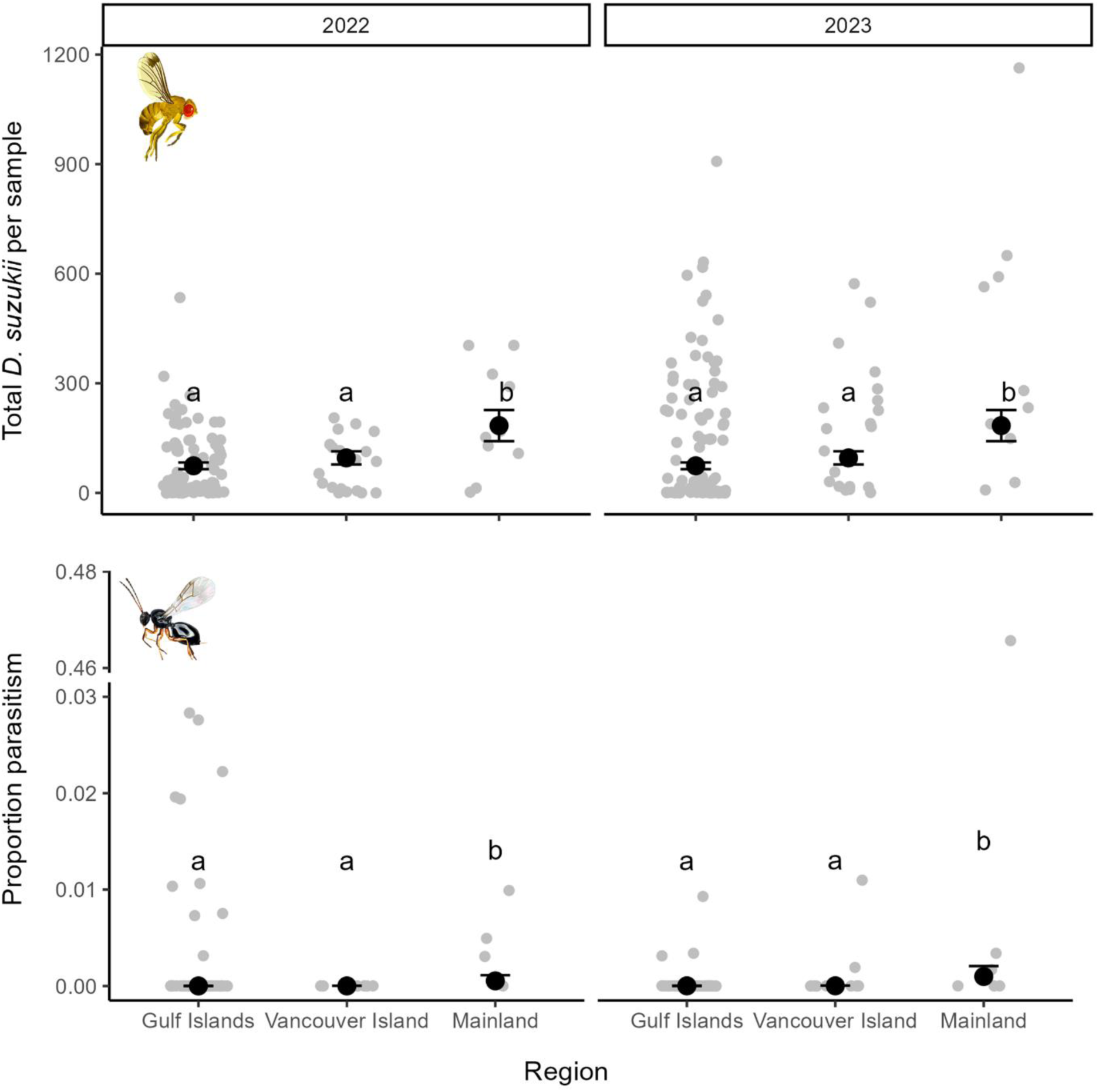
The number of *Drosophila suzukii* per sample of 45 *Rubus armeniacus* berries and the proportion of *D. suzukii* larvae parasitized in the three sampled regions in each of the two years of the study. Each grey point represents the raw data from each sampling event (site/year/date combination). Black points show estimated marginal means (±SE) from fitted mixed models. Multiple comparisons were done with the Sidak method; estimates with labels not containing the same letter are statistically different (p < 0.05). Note the axis break in the lower panel to show an outlier datapoint.

A total of 277 parasitoids (223 *L. japonica* and 44 *G. kimorum*) emerged from berry samples. Parasitoids were present (found in at least one of the four sampling events over the two-year study period) at all five mainland sites, 3/10 sites on Vancouver Island, and 11/43 sites on the Gulf Islands. Both *L. japonica* and *G. kimorum* were present at sites on the mainland and Vancouver Island, whereas we only found *L. japonica* on the Gulf Islands. We detected *L. japonica* at least once on five Gulf Islands (Cortes, Quadra, Hornby, Gabriola, Salt Spring), but only detected the parasitoid in both years on three of these islands (Cortes, Quadra, and Hornby).

The proportion of parasitized *D. suzukii* varied among regions (χ²_2_=10.25; p = 0.0059) and tended to be higher on the mainland than Vancouver Island and the Gulf Islands (Figure 3). Proportion parasitism per collecting event varied from 0 to 0.47 but was very low overall (<1% on average) in all three regions (Figure 3). Parasitism was higher in 2022 than 2023 (χ²_1_=11.16; p < 0.001) and varied over the course of the sampling period (χ²_3_=156.49; p < 0.0001). There was no residual spatial autocorrelation in proportion parasitism (Moran’s *I* = -0.026, p = 0.59).

### Variation in density of D. suzukii among the twelve Gulf Islands

On the Gulf Islands, *D. suzukii* densities varied by island (GLMM; χ²_11_=84.27; p < 0.0001) but were similar between the two years of the study (χ²_1_=0.46; p =0.50). Over the course of the sampling period, *D. suzukii* densities on islands started low, increased strongly, and then declined somewhat at the end of the sampling period (χ²_3_=184.78; p < 0.0001) (Figure S4). There was no residual spatial autocorrelation in *D. suzukii* densities on the Gulf Islands after accounting for these variables (Moran’s *I* = -0.053, p = 0.71).

### Associations between island characteristics and parasitoid presence

*Leptopilina japonica*, the only species of parasitoid reared from *D. suzukii* on the Gulf Islands, was marginally more likely to be detected on islands with higher average *D. suzukii* densities (GLM; χ²_1_ = 4.61, p = 0.032) and larger islands (χ²_1_ = 4.61, p = 0.066) (Figures 4, 5). The presence of *L. japonica*, however, was clearly not associated with the amount of ferry traffic to each island (χ²_1_ = 0.15, p = 0.70) (Figure 4). We did not detect spatial autocorrelation in the residuals of the GLM containing *D. suzukii* density and island size (Moran’s *I* = -0.036, p = 0.34). *Leptopilina japonica* was not any more likely to be found on islands with a greater number of sampling sites (χ²_1_ = 0.14, p = 0.71).

**Figure 4.**
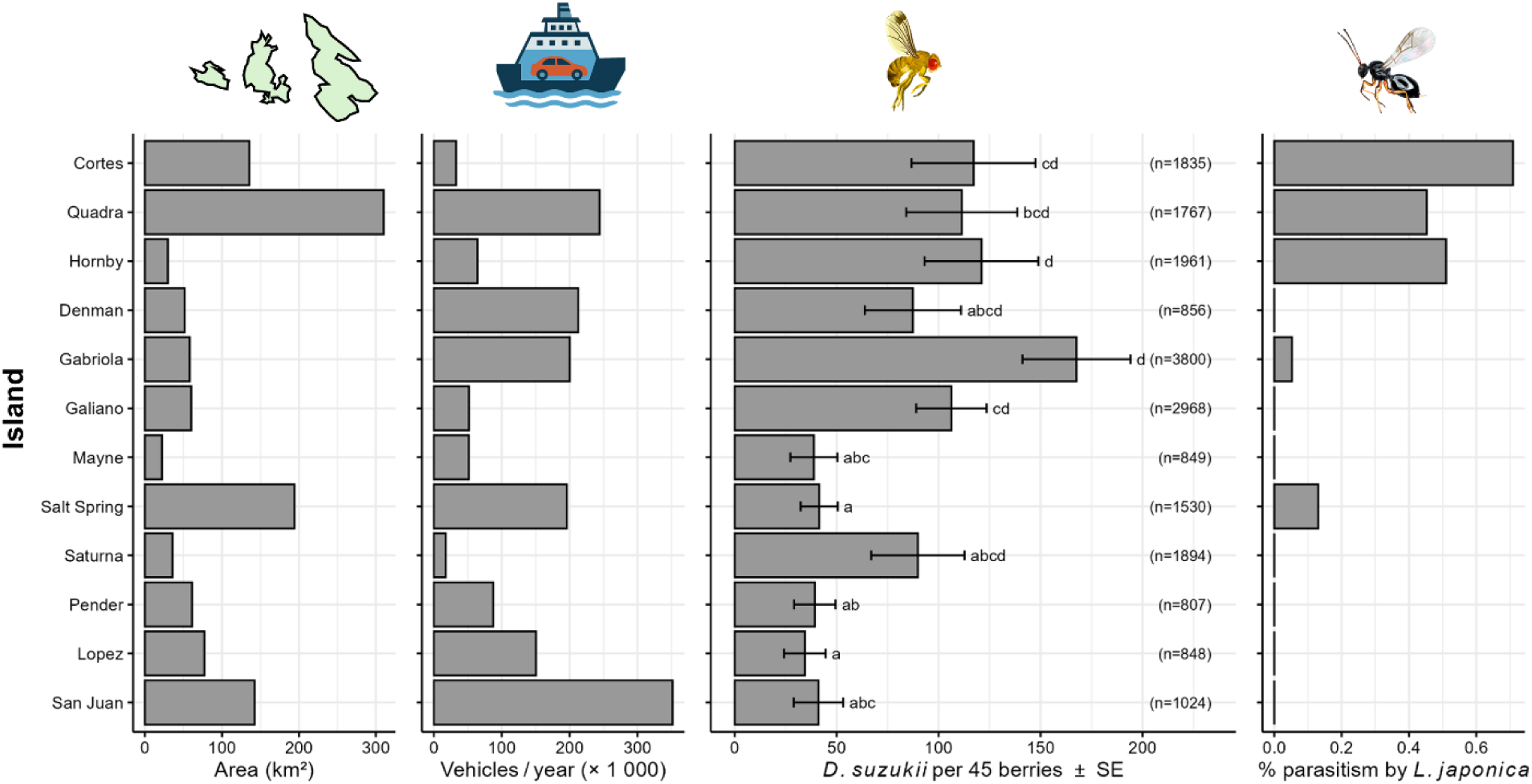
Metrics for each of the Gulf Islands sampled in the study (arranged from highest to lowest latitude from top to bottom): area, ferry vehicle traffic, marginal mean (± SE) number of *Drosophila suzukii* per sample of 45 *Rubus armeniacus* berries (estimated from a generalized linear mixed model; see Results – Statistical analyses) with the total number of *D. suzukii* shown in parentheses, and the percentage of all *D. suzukii* collected that were parasitized by *L. japonica*. In the third panel, letters are from Tukey multiple comparisons. Means labeled with different letters are different (p < 0.05).

**Figure 5.**
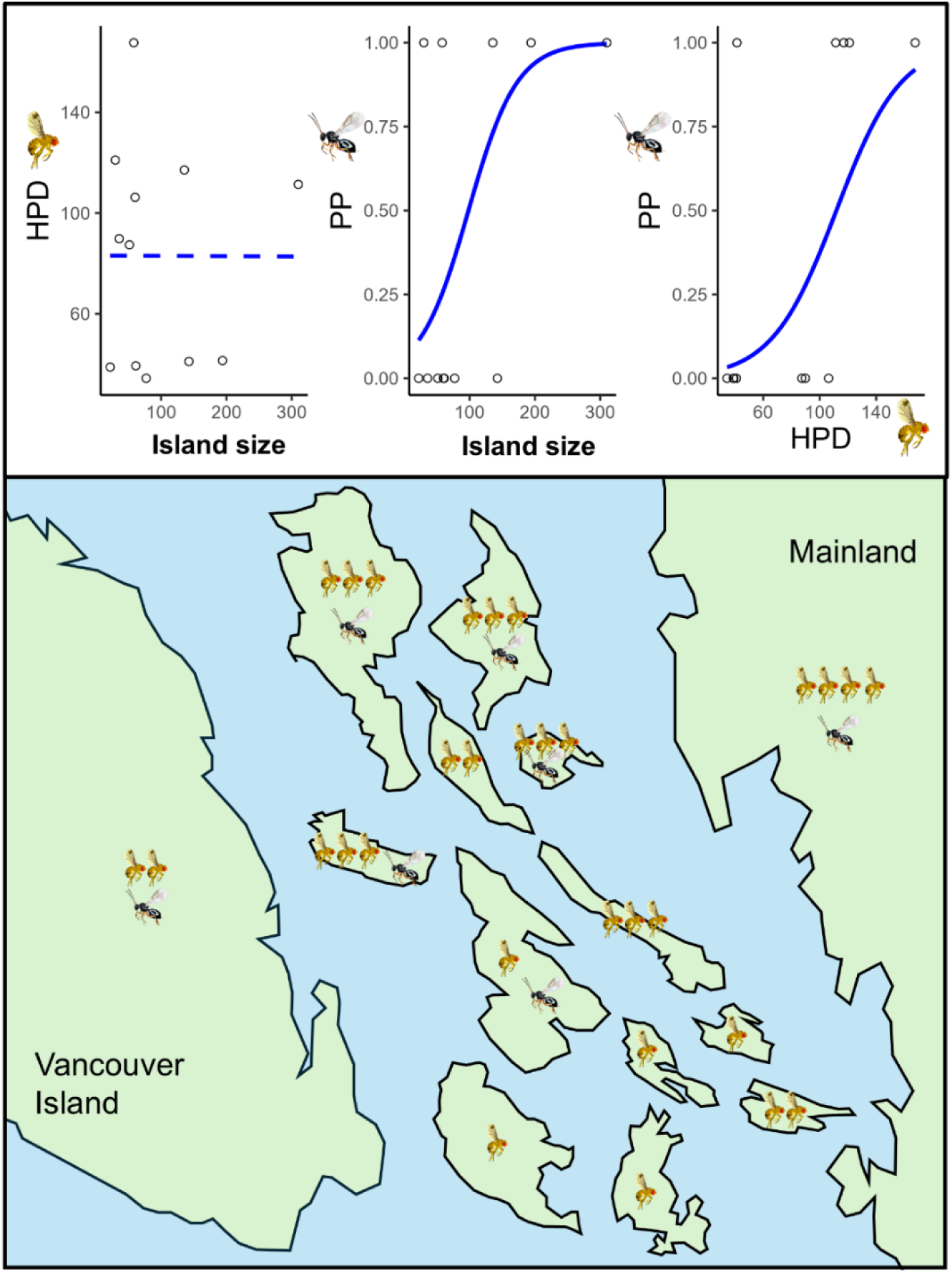
An illustrative summary of the results of the study compared against the framework presented in Figure 1. The upper panel shows marginal effects plots of the relationships detected between mean *D. suzukii* population density (HPD), parasitoid presence (PP), and island size (km^2^) (See Results for statistical details). Solid lines indicate significant (p < 0.05) or marginal relationships (0.05 < p < 0.07) whereas dotted lines show non-significant (p > 0.30). The lower panel shows the relative density of *D. suzukii* throughout the study area (indicated by the relative number of flies), the presence of parasitoids (indicated by wasps), and the relative size of the Gulf islands (which is to scale). The position of the Gulf Islands relative to each other and the scale, shape and position of Vancouver Island and the Mainland are for illustration only, and are not accurate (see Figure 2 for actual geographic layout).

The positive association between island area and parasitoid presence was not mediated by *D. suzukii* abundance. On its own, island area was not a strong predictor of parasitoid presence (β = 0.011 ± 0.008; χ²₁ = 2.53; p = 0.13). When controlling for *D. suzukii* abundance, the effect of island area became stronger (β = 0.027 ± 0.016; χ²₁ = 4.61; p = 0.066).

Island area was positively correlated with ferry vehicle traffic (*r* = 0.53) (Figure S3). The effect of island area was nearly identical whether or not ferry traffic was included in the model with *D. suzukii* abundance (β = 0.027 ± 0.016 without ferry traffic; β = 0.026 ± 0.015 with ferry traffic). By contrast, ferry vehicle traffic itself was not an important predictor of parasitoid presence when included alongside *D. suzukii* abundance (χ²₁ = 0.44; p = 0.51).

*Drosophila suzukii* density tended to be higher on islands where parasitoids were present (LM; F_1_ = 6.55, p = 0.034), but there was no association between *D. suzukii* density and island area (F_1_ = 1.61, p = 0.24), nor was there an effect of island area’s interaction with parasitoid presence (F_2_ = 0.02, p = 0.89).

## Discussion

Our study of relatively large continental islands with high levels of human activity in the Pacific Northwest of North America revealed that an invasive insect pest, *D. suzukii*, is essentially ubiquitous on both island and mainland areas over a large geographic area less than 20 years after its initial detection. Its accidentally introduced parasitoid wasps, while present on the mainland and Vancouver Island (as previously known; Abram et al. 2020, 2022b), were only present on five of the twelve Gulf islands. Only one out of two parasitoid species (*L. japonica*) appears to have established on any of the Gulf Islands we surveyed. These parasitoids were not most common on islands with higher levels of human activity – rather, they tended to be found on larger islands with higher *D. suzukii* populations, including two relatively remote, northern islands (Cortes and Quadra Islands). Below, we discuss these results in the context of each of our original predictions.

Our first prediction, consistent with island biogeography theory, was that parasitoids would be more likely to establish on larger islands because the larger area would provide a larger and more stable resource base for these parasitoids (Schoener 1989; Holt et al. 1999; Holt 2009), which occupy a relatively high trophic level and depend on certain drosophilid hosts (and their basal food resources, fruiting plants) to persist. We found some support for this prediction – *L. japonica* was marginally more likely to be present on larger islands, possibly indicating that larger area may help to stabilize host-parasitoid interactions (Hassell et al. 1991; Holt 1999; Hassell 2000; Holt 2009), may provide a larger overall resource base for the parasitoid (Schoener 1989), or may simply favor the persistence of the parasitoid through simple numerical effects (Srivastava et al. 2008). Larger islands may support higher plant diversity (Kohn and Walsh 1994), providing early-fruiting hosts that sustain *D. suzukii* and allow *Leptopilina japonica*— which has at least two generations per year in British Columbia (Abram et al. 2022a; Rossi Stacconi et al. 2025)—to complete its first generation before *Rubus armeniacus* fruits in mid-July (Abram et al. 2022a). However, the effect of island size was marginal – parasitoids were absent on one of the largest islands we surveyed (San Juan), and detected in only one of two years on the largest island (Salt Spring). They were also found on small to intermediate-sized islands as well (Gabriola, Hornby), suggesting that other factors are at play. For example, smaller islands may be able to support parasitoid establishment if their host densities are high enough (see below) – indeed, the two smallest islands where parasitoids were detected also had the highest mean *D. suzukii* densities. It is also possible that the limited variation of island sizes we sampled prevented the detection of a stronger relationship between island size and parasitoid presence. That is, the largest islands we surveyed may have been near the lower threshold for parasitoid establishment, or we did not survey small enough islands where parasitoids would more consistently be absent.

Our second prediction was that parasitoids of *D. suzukii* would be more likely to establish on less isolated islands. In island landscapes with relatively high levels of human activity, we would have expected variation in transport of goods and people (here, estimated by the amount of vehicles arriving by ferry to each island) to be the main source of differences in island isolation (Helmus et al. 2014). Surprisingly, we found no evidence that parasitoids were more likely to be present on islands where more vehicles arrive by ferry each year or where there are more year-round human residents. Parasitoids were present both on sparsely populated islands with relatively low ferry traffic as well as islands with relatively high populations and hundreds of thousands of vehicles arriving per year. It is possible that we simply used the wrong metric of isolation, that natural dispersal mechanisms are more important, and that geographic distance from mainland areas where the parasitoids are known to be common would be a better metric of isolation. However, our data tend in the opposite direction – parasitoids were most likely to be consistently present on three of the most northern islands most isolated from the mainland area of British Columbia and Washington where the parasitoids were previously known to be common (Abram et al. 2020, 2022b; Beers et al. 2022; Appendix S1: Methods S1; Results S1). A perhaps more likely explanation is that all the islands we surveyed, which each receive a minimum of ∼17,000 vehicles per year, receive enough human traffic to introduce the parasitoids (i.e., the propagule pressure is high enough to introduce them everywhere), but that other factors such as island size (see above), host population densities (see below) and potentially climatic suitability (see below) determine whether or not they establish breeding populations after they are introduced.

Our third prediction was that if the parasitoids are in their early phases of establishment, their presence would be positively associated with average host densities, because islands with higher host populations (which we might expect to be the largest islands; Connor et al. 2000) would be more receptive to their establishment. If the parasitoids were found to be well-established and strongly reducing the abundance of host populations (as per the enemy release hypothesis; Keane and Crawley 2002), we might expect them to be more likely to be found where host densities are relatively low. More consistent with the first scenario, we found that parasitoids were more likely to be found on islands where host densities were higher, but contrary to our predictions, host densities were not associated with island size. Parasitism levels were very low throughout the study region overall relative to past studies (Abram et al. 2020; Abram et al. 2022a), but especially on islands, indicating that parasitoids may be in their early phase of establishment on these islands. The reasons for the very low parasitism levels during this time period are not known conclusively, but are suspected to have been at least partially due to the negative effect of the 2021 western North American ‘heat dome’ extreme heat event (Zhang et al. 2023) on insect populations in the year preceding the beginning of the study. Because parasitism levels were so low, and may have sometimes been just above or below our sampling regime’s ability to detect parasitism, our absence/presence designations may be better viewed as detection probabilities given our sampling regime. Regardless, there is not currently any evidence from our data that *L. japonica* is exerting strong population control of *D. suzukii* on these islands – if it were, we would have expected the per-berry density of *D. suzukii* to be lower on islands where the parasitoids were present, which they were not. In fact, regionally, *D. suzukii* were the most abundant on islands where parasitoids were present, and even more abundant on the mainland, where parasitism levels were highest and where the parasitoids are known to have been present for at least 3-6 years (Abram et al. 2020). These findings suggest that at least in our study’s timeframe, enemy release and biological control are not the dominant factors determining variation in *D. suzukii* abundance on islands or in our study region as a whole. It is possible, however, that this could change if parasitoid establishment on islands progresses in the coming years. In addition, if population dynamics are cyclical (e.g. Myers and Cory 2013), longer-term datasets may be needed to detect signals of top-down population regulation of *D. suzukii* by its parasitoids.

An important caveat in interpreting the relationship between *D. suzukii* densities, island size, and parasitoid presence is that our estimate of relative *D. suzukii* densities (number of larvae per berry) is not the same as total abundance (which may be more important for longer-term parasitoid establishment), as it does not consider variation in the total amount of *R. armeniacus* and other host plants on each island. Total *D. suzukii* abundance on an island depends on both larval density per fruit and the total number of fruits available. More detailed surveys of fruiting plant cover and diversity were not possible given the resources available to this study (see Methods), and would be needed to better estimate the population size of *D. suzukii*, and to more conclusively tease out the importance of host density *versus* abundance for parasitoid establishment. However, given that most islands had broadly similar land cover composition (Appendix S1: Figure S1), and most of the land cover types on the islands would be expected to host *R. armeniacus*, it may be reasonable to generally assume that total resource availability for *D. suzukii* scales positively with area, and larger islands also have higher total *D. suzukii* abundances. Thus, at least some of the modest effect of island size on parasitoid presence that we detected could have been mediated by island size’s likely positive effect on total *D. suzukii* abundance (independent of density).

Our final prediction was that the less specialized species (*L. japonica*) would be more likely to establish on an island with a given set of characteristics than the more specialized species (*G. kimorum*). This prediction was based on previous hypotheses (Freeland 1983) and empirical studies (Steffan-Dewenter and Tscharntke 2000; Kruess and Tscharntke 2000) suggesting that specialist parasitoids and parasites are more vulnerable to demographic stochasticity (which is higher on smaller islands) of the hosts on which they are specialized. This was indeed the case: both parasitoid species were found on the mainland and Vancouver Island, while only *L. japonica* was detected on the Gulf Islands. *Leptopilina japonica* is able to parasitize other species of Drosophilidae (e.g. in the *obscura* and *melanogaster* species groups) that are known to be common and abundant in our study region and develop on a variety of substrates other than fresh fruit (Rossi Stacconi et al. 2025) at times of the season when *D. suzukii* is less abundant (P.K. Abram, personal observations). This may help reduce this parasitoid’s trophic dependency on *D. suzukii*, which reproduces mostly on fresh fruits and so is reliant on the availability and seasonal phenology of fruiting plants. It is possible that *G. kimorum* is unable to establish on islands because – especially in the early summer when the first parasitoid generation emerges from overwintering – *D. suzukii* populations are at very low densities (more so on islands; Figure 3) and due to its high level of specialization on *D. suzukii* inside ripening fruit (Stahl et al. 2024), *G. kimorum* cannot ‘fall back’ on other alternative Drosophilidae host species that are present at that time.

We noted an unexpected trend: the parasitoid *L. japonica* appeared to be more likely to be present on higher-latitude islands, which also tended to have higher *D. suzukii* abundances (Figure 3; Appendix S1: Results S1; Figure S3). One possible explanation is that more southerly islands (San Juan, Lopez) are considerably cooler than more northerly islands during the seasonal period where *D. suzukii* and its parasitoids reproduce and develop (Figure S2), potentially allowing populations of *D. suzukii* on more northerly islands to build to higher levels each season, in turn making parasitoid establishment more likely. This anomalous temperature-latitude trend in our study region is likely due to regional variation in ocean temperatures and wind patterns (Ruping Mo, Environment and Climate Change Canada, personal communication), and may create a unique landscape where island characteristics and counterintuitive regional climatic variation may interact to influence invasion and consumer-resource dynamics.

This study system and region constitutes a ‘natural experiment’ in biological control and island biogeography involving invasions by a non-native insect and its parasitoid wasps, in an island landscape highly shaped by human activity. While our two-year study presented here is clearly just a starting point, we have succeeded in capturing a snapshot early in the progression of the parasitoids’ invasion of these relatively large islands with high levels of human activity. The ubiquity of the nonnative *D. suzukii* in our study region is a particularly striking result, which could in part be because the non-native plant *R. armeniacus* is so abundant and widespread. This could provide further support to the idea that the spread of non-native plants provides opportunities for subsequent invasions by their insect herbivores (Bertelsmeier et al. 2024) – including on islands that might otherwise be unable to support the invading herbivores. An extension of this idea, articulated in the ‘receptive bridgehead hypothesis’ (Weber et al. 2021) is that the arrival of these nonnative herbivores and the high levels of abundance they reach may in turn facilitate the establishment of their natural enemies that occupy higher trophic levels. Our study, finding that islands with higher host densities are more likely to result in non-native parasitoid establishment, supports this hypothesis. It is still unclear whether the establishment of these parasitoids on larger islands with higher host densities will affect the relationship between island characteristics and invasive pest densities in the long term, as per our trophic dampening hypothesis. This sets the stage for using this study system and region to investigate how biological control operates in island landscapes in the anthropocene.

## Acknowledgements

We thank Yonathan Uriel, Jessie Moon, Victoria Makovetski, Judy Myers, Julia Gray, Adriana Barsan, and Claire Winslow for help with travel arrangements, sample collection, and data collection. We thank Aysha McConkey for creating the parasitoid and fly artwork used in some of the figures. Thanks to Alex Cannon and Ruping Mo (Environment and Climate Change Canada) for help interpreting weather data. This work was supported by funding to PKA and MTF from Agriculture and Agri-Food Canada (projects J-003401, J-002839, J-002646). We acknowledge the support of the Natural Sciences and Engineering Research Council of Canada (NSERC) Alliance Missions Program, ALLRP 570736-2021 to JC. We thank Diane Srivastava, Rachel Germain, and Ian Kaplan for helpful comments on an earlier version of the manuscript. Artificial Intelligence (ChatGPT versions 4o and 5) was used to assist with data formatting and the production of figures in R, as well as to produce the drawing of the ferry in Figure 4. The authors take full responsibility for the final R code and figures.

## Author contributions

PKA and JC led the conceptualization of the study. All authors contributed to planning the data collection. PKA, JT, LBN, BD, MTF and CCC collected and processed samples. PKA wrote the first draft of the manuscript and conducted the data analysis with support from JC. All authors read, edited and approved the manuscript.

## Notes

### Competing Interest Statement

The authors have declared no competing interest.

