## Appendix S1 for "Island invasions by the non-native vinegar fly *Drosophila suzukii* and its parasitoid wasps"

For the pre-print:

**Island invasions by the non-native vinegar fly *Drosophila suzukii* and its parasitoid wasps**

Paul K. Abram, Elizabeth H. Beers, Charlie C. Coslor, Benjamin Diehl, Michelle T. Franklin, Louis B. Nottingham, Jason Thiessen, Matthew Tsuruda, Juli Carrillo

**Methods S1**

*Association between latitude and parasitoid presence*

Upon inspecting our results, we noted that parasitoids seemed to be more likely to be found on islands at higher latitudes (Main Text: Figure 4). We tested for the statistical significance of this association using a GLM with Firth's bias-reduced penalized maximum likelihood estimation (see Main Text: Methods for details) with parasitoid presence as the response variable. Latitude was the sole predictor variable included in the model; when it was added to a model already containing island size and mean *D. suzukii* density, the model failed to converge, probably due to its relatively strong correlation with *D. suzukii* density (Figure S3) and the low number of residual degrees of freedom in the model containing three predictor variables.

*Land cover analyses*

We quantified land cover composition around each study site, and on each of the 12 Gulf Islands as a whole, using the North American Land Change Monitoring System (NALCMS) 30 m resolution land cover dataset for 2020 (CEC 2023). For the site-level analysis, raster layers were clipped to circular buffers with a 0.75 km radius around each site. Zonal histograms in QGIS (version 3.16) were used to calculate the proportion of pixels assigned to each of the 19 NALCMS land cover classes around sites and on whole islands. These values were exported, summarized and analyzed in R version 4.4.1 (R Core Team 2024). Based on existing literature showing the *D. suzukii* and parasitoid abundance or density is often (although not always) positively related to percent forest cover (Pelton et al. 2016; Cahenzli et al. 2018; Haro-Barchin et al. 2018; Santoiemma et al. 2018; Tonina et al. 2018; Hogg and Daane 2024), we selected percent forest cover as the focal habitat composition metric for subsequent analyses. Forest cover was defined as the sum of pixels classified as forest types (temperate or sub-polar needleleaf, temperate or sub-polar broadleaf deciduous, and mixed forest). Percent forest cover was calculated relative to the total number of pixels within each buffer. For the site-level analysis, as several sampling sites were close to coastlines, we included ocean in the calculation of percent forest cover.

We first compared percent forest cover at the site level among the three region types (Gulf Islands, Vancouver Island, and the mainland) using a GLM with quasi-binomial error distribution (after detecting overdispersion in the original fitted model) and an F-test for significance testing. Then, to test whether percent forest cover explained additional variation in *D. suzukii* densities or proportion parasitism at each site, we added it as a fixed factor to the simplified regional GLMMs (see Main Text: Methods for details) containing region (Mainland, Vancouver Island, or the Gulf Islands) and Julian date (non-linear spline, df=3) as fixed factors and site nested within region type as a random factor. We then used Type II Wald chi-squared tests to test the statistical significance of percent forest cover as a predictor of *D. suzukii* density and proportion parasitism within these statistical models. Finally, we tested whether percent forest cover at the island level was associated with *L. japonica* presence using a GLM with Firth's bias-reduced penalized maximum likelihood estimation (see Main Text: Methods for details). It was tested both as a sole predictor variable as well as a predictor added to a model already containing island area and *D. suzukii* density.

*Distance of each island to mainland populations of parasitoids*

We wanted to know whether the geographic distance of each island to the mainland might be associated with parasitoid presence, to justify the fact that was not used as the hypothesized driver of island isolation *a priori*. Defining the distance of each island in our geographic study area to the mainland was challenging, especially given that many mainland areas have not been surveyed for the presence of *D. suzukii* parasitoids. However, as a best approximation, we defined the distance to the mainland of each island as the shortest linear distance between the midpoint of the sampling sites on the island and a known mainland (British Columbia, Canada or Washington State, USA) site where *L. japonica* is present based on past studies (Abram et al. 2020; Abram et al. 2022; Beers et al. 2022; Czokajlo et al. 2025) as well as new detections in coastal mainland areas in the current study. Given that Vancouver Island can be considered a 'second mainland', we conducted a second analysis where the distance to the mainland was defined as the shortest distance between the midpoint of sampling sites on each island and a mainland *or* Vancouver Island site where *L. japonica* is known to be present. Two separate GLMs (with Firth's bias-reduced penalized maximum likelihood estimation; see Main Text: Methods for details) with parasitoid presence as a predictor variable and distance to the mainland (either excluding or excluding Vancouver Island) were run. Each of the distance to the mainland metrics were the sole predictor variable added to each of the two models; when they were added to models already containing island size and mean *D. suzukii* density, the models failed to converge.

**Results S1**

*Association between latitude and parasitoid presence*

Parasitoids were more likely to be detected on islands at higher latitudes (GLM; $\chi^2_1 = 4.51$, p = 0.034). Latitude was also positively correlated with marginal mean *D. suzukii* density (Pearson's $r = 0.64$; p = 0.026).

*Land cover analyses*

Percent forest cover around each sampling site varied among the three study regions (GLM; $F_{2,55} = 35.07$, $p < 0.0001$), with the Gulf Island sampling sites having higher percent forest cover in the area surrounding them than Vancouver Island or mainland sites (Figure S1). Percent forest cover was not associated with *D. suzukii* density (GLMM; $\chi^2_1 = 1.04$, p = 0.31) or the proportion of *D. suzukii* that were parasitized (GLMM; $\chi^2_1 = 0.27$, p = 0.60). The presence of parasitoids on islands was not associated with the islands' percent forest cover either as a sole predictor variable (GLM; $\chi^2_1 = 0.36$, p = 0.55) or when added to a model containing island area and *D. suzukii* density (GLM; $\chi^2_1 = 0.68$, p = 0.41).

*Distance of each island to mainland populations of parasitoids*

Parasitoids (*L. japonica*) were marginally more likely to be detected on islands that were further from known mainland populations of *L. japonica* (GLM; $\chi^2_1 = 3.29$, p = 0.060). Notably, the distance of islands to mainland populations of *L. japonica* was highly correlated with latitude (Pearson's $r = 0.97$, $p < 0.0001$). There was no association between *L. japonica* presence on islands and the distance to the closest Vancouver Island or Mainland population (GLM; $\chi^2_1$ = 2.30, $p = 0.13$).

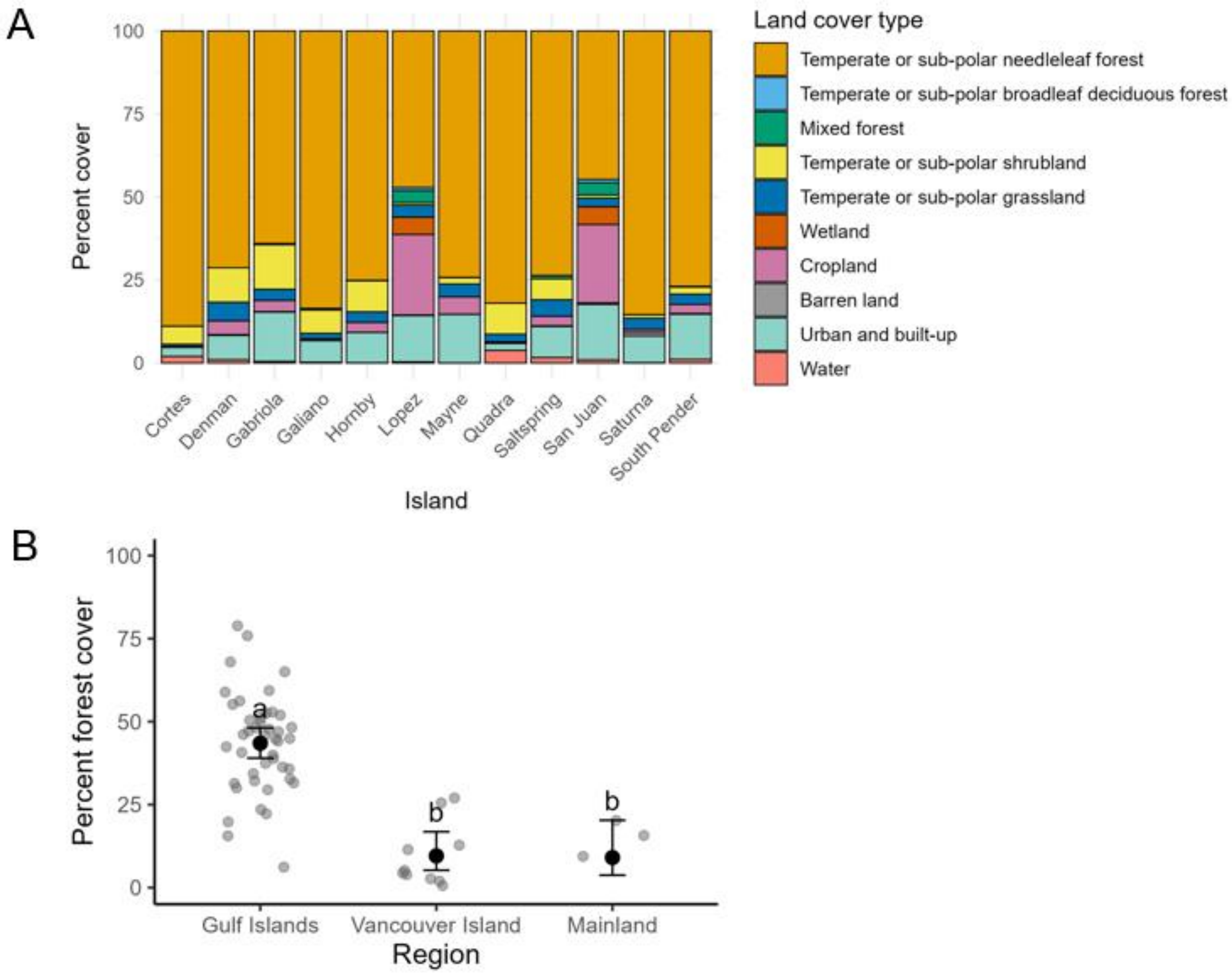


**Figure S1.** Land cover characteristics of the 12 Gulf Islands and sampling sites in each of three regions sampled in this study (data from CEC 2023). (A) The percentage of each type of land cover of each of the 12 Gulf Islands; (B) The percentage of forest cover in a 750m radius around each sampling site in each of the three main study regions. Each grey point represents one sampling site. Black points with error bars show estimated marginal means (±SE) from a fitted GLM. Multiple comparisons were done with the Tukey method; estimates with labels not containing the same letter are statistically different ($p < 0.05$).

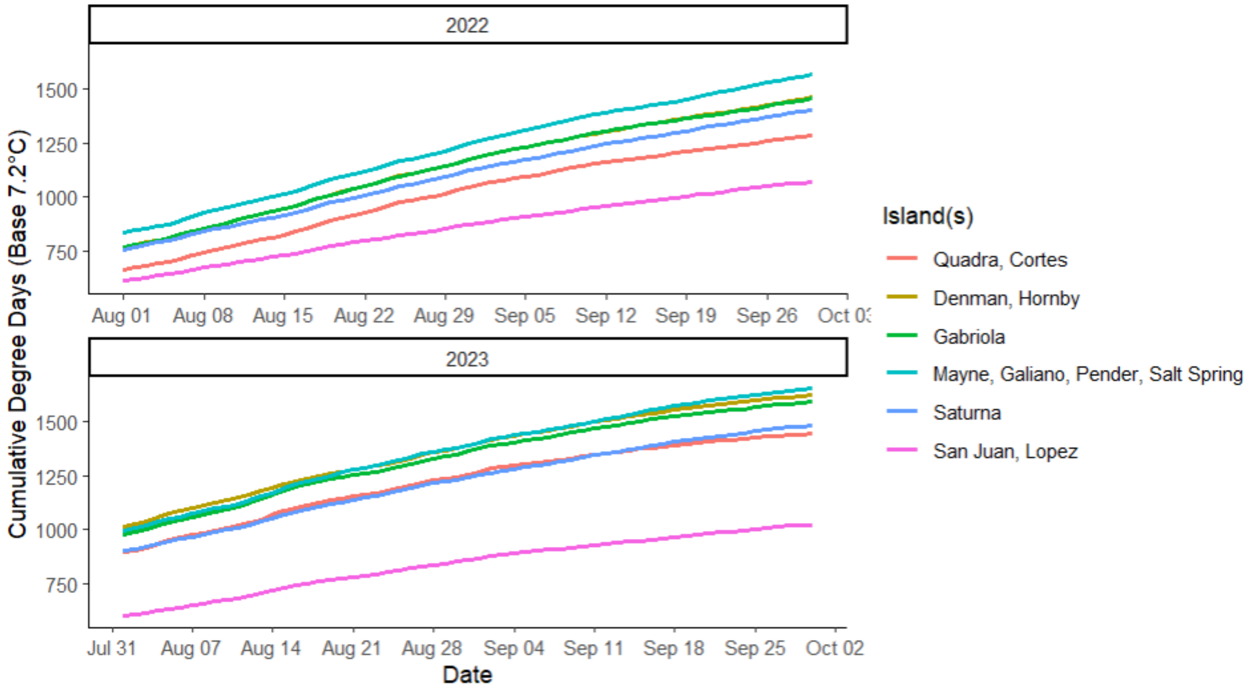


**Figure S2.** The number of heat units (degree days; base temperature: 7.2°C) that accumulated over the course of the season on each island in each year of the study. Islands (grouped by nearest weather station) are ordered from highest to lowest latitude in the legend. Degree days were calculated from weather data from stations as close as possible to each sampled island (Environment and Climate Change Canada 2024; National Weather Service 2024).

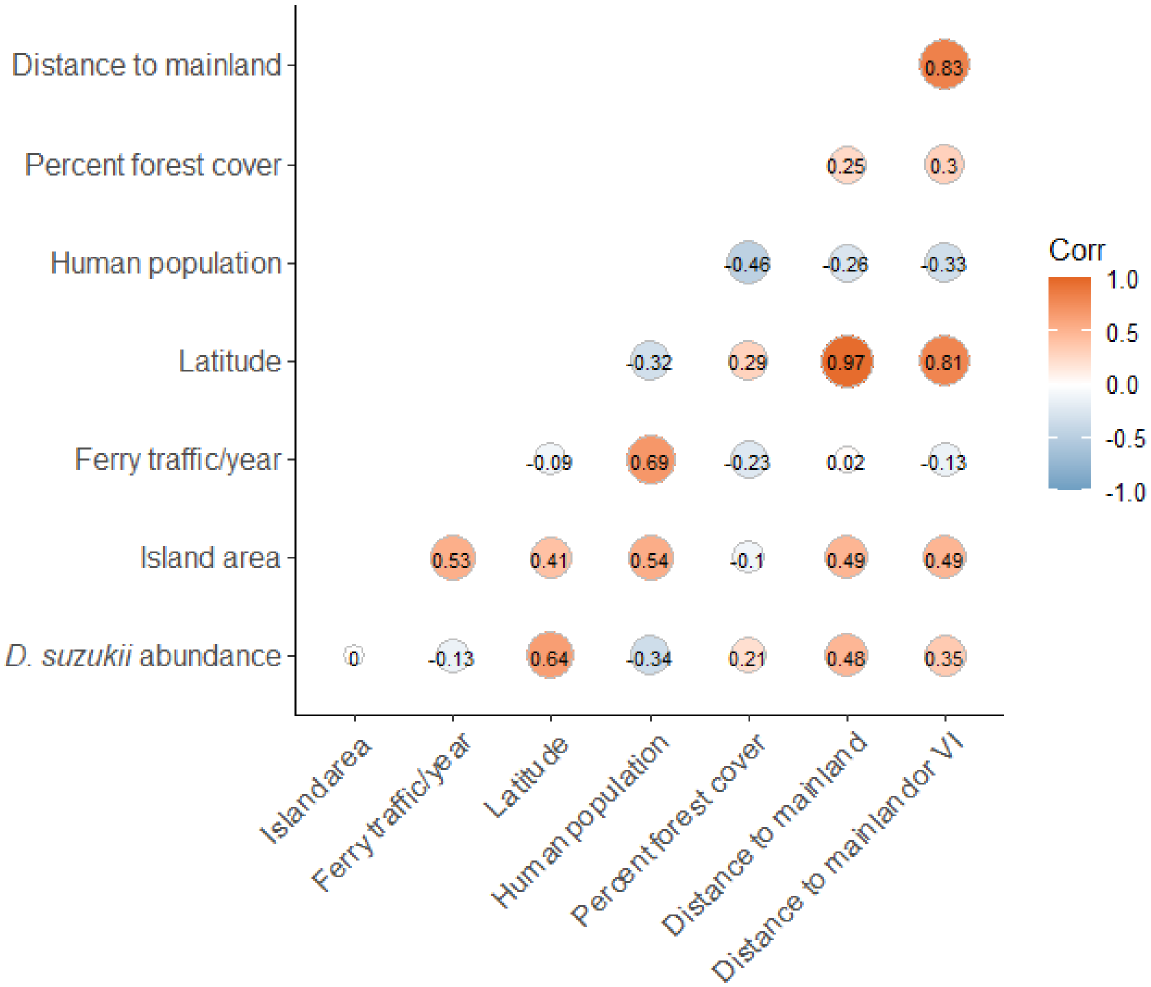


**Figure S3.** Correlation matrix showing correlations (Corr – Pearson's correlation coefficient) among the main explanatory factors as well as the variables of interest in the post-hoc analyses for the 12 Gulf Islands in this study. VI – Vancouver Island.

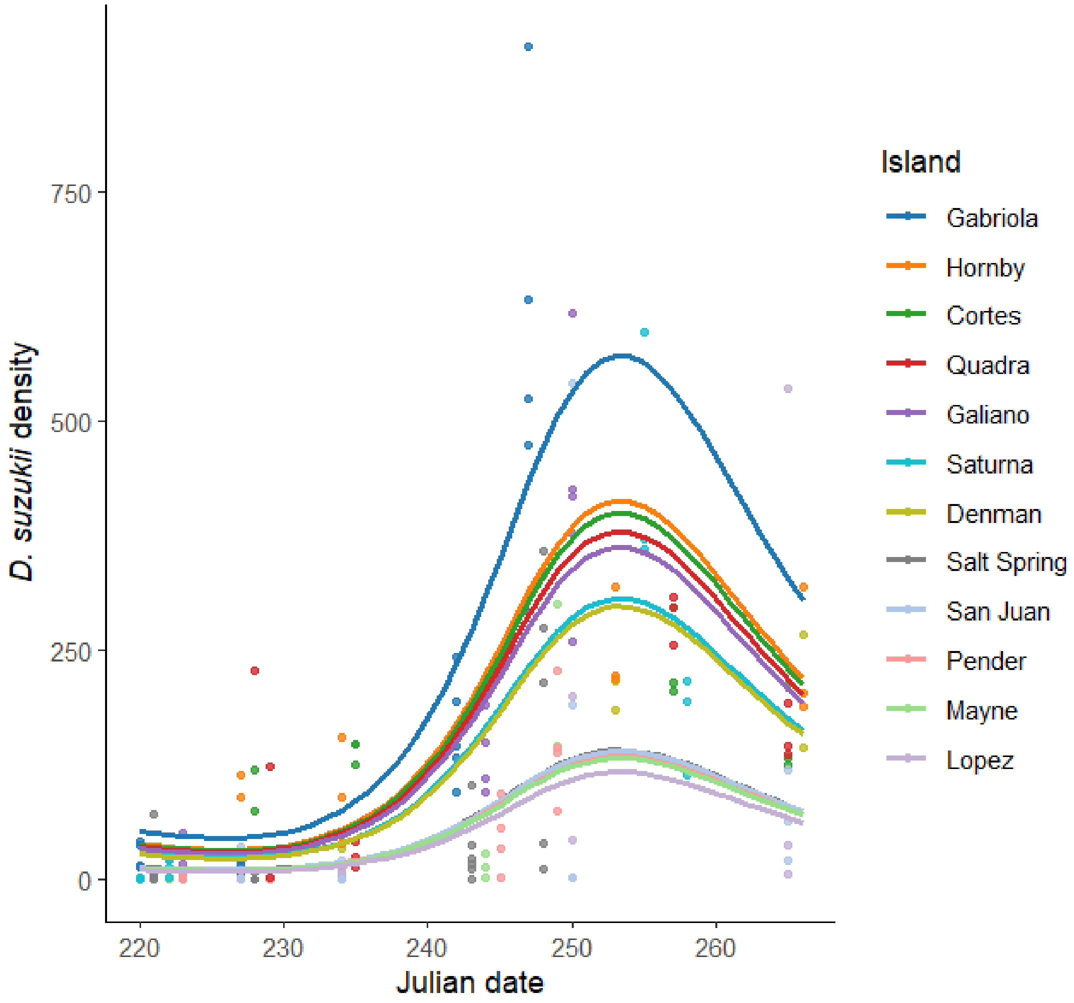


**Figure S4.** Trends in *Drosophila suzukii* densities (number emerging per sample of 45 berries) over the study's sampling period (Julian date; both years combined) on each island. Points show observed densities, while lines show predictions from a generalized linear mixed model (see Results, Variation in density of *D. suzukii* among the twelve Gulf Islands).

**References S1**

Abram, P. K., A. E. McPherson, R. R. Kula, T. Hueppelsheuser, J. Thiessen, S. J. Perlman, C. I. Curtis, J. L. Fraser, J. Tam, J. Carrillo, M. W. Gates, S. Scheffer, M. Lewis, and M. L. Buffington. 2020. "New Records of *Leptopilina*, *Ganaspis*, and *Asobara* Species Associated with *Drosophila suzukii* in North America, Including Detections of *L. japonica* and *G. brasiliensis*." *Journal of Hymenoptera Research* 78: 1–17.

Abram, P. K., M. T. Franklin, T. Hueppelsheuser, J. Carrillo, E. Grove, P. Eraso, S. Acheampong, L. Keery, P. Girod, M. Tsuruda, M. Clausen, M. L. Buffington, and C. E. Moffat. 2022a. "Adventive Larval Parasitoids Reconstruct Their Close Association with Spotted-Wing Drosophila in the Invaded North American Range." *Environmental Entomology* 51(4): 670–678.

Czokajlo, R., C. Looney, L. Nottingham, B. Diehl, P. Abram, T. Northfield, P. Smytheman, and E. Beers. In press. "Distribution of Three Figitid Parasitoids of Drosophilidae in Washington State: A Tale of Two Ecozones." *Journal of Economic Entomology*.

Cahenzli, F., I. Bühlmann, C. Daniel, and J. Fahrentrapp. 2018. "The Distance between Forests and Crops Affects the Abundance of *Drosophila suzukii* during Fruit Ripening, but Not during Harvest." *Environmental Entomology* 47: 1274–1279.

Commission for Environmental Cooperation (CEC). 2023. "2020 North American Land Cover 30-Meter Dataset." North American Land Change Monitoring System. https://www.cec.org/north-american-environmental-atlas/land-cover-30m-2020/.

Environment and Climate Change Canada. 2024. *Historical Climate Data*. https://climate.weather.gc.ca

Haro-Barchin, E., J. Scheper, C. Ganuza, G. A. de Groot, F. Colombari, R. van Kats, and D. Kleijn. 2018. "Landscape-Scale Forest Cover Increases the Abundance of *Drosophila suzukii* and Parasitoid Wasps." *Basic and Applied Ecology* 31: 33–43.

Hogg, B. N., and K. M. Daane. 2024. "Landscape Effects on Seasonal Abundance of *Drosophila suzukii* and Its Parasitoids in California Caneberry Fields." *Agricultural and Forest Entomology* 27: 304–315.

National Weather Service. 2024. *Historical Weather Data for Washington State*. U.S. National Oceanic and Atmospheric Administration. https://www.weather.gov

Pelton, E., C. Gratton, R. Isaacs, S. Van Timmeren, A. Blanton, and C. Guédot. 2016. “Earlier Activity of *Drosophila suzukii* in High Woodland Landscapes but Relative Abundance Is Unaffected.” *Journal of Pest Science* 89: 725–733.

R Core Team. 2024. *R: A Language and Environment for Statistical Computing.* Vienna: R Foundation for Statistical Computing. https://www.R-project.org

Santoiemma, G., N. Mori, L. Tonina, and L. Marini. 2018. “Semi-Natural Habitats Boost *Drosophila suzukii* Populations and Crop Damage in Sweet Cherry.” *Agriculture, Ecosystems & Environment* 257: 152–158.

Tonina, L., N. Mori, M. Sancassani, P. Dall’Ara, and L. Marini. 2018. “Spillover of *Drosophila suzukii* between Non-Crop and Crop Areas: Implications for Pest Management.” *Agricultural and Forest Entomology* 20: 575–581.
